# Gravid *Anopheles stephensi* Detects Indole for Oviposition Despite Ablation of Antennae and Maxillary Palps

**DOI:** 10.64898/2026.01.19.700218

**Authors:** John Agbetsi, Jiannong Xu

## Abstract

Oviposition site selection is critical for mosquito population dynamics. Gravid mosquitoes rely on chemical cues to identify suitable breeding habitats. However, the sensory mechanisms governing this behavior in *Anopheles stephensi* remain poorly understood. Here, we examined the role of indole, a microbial volatile associated with aquatic environments, in oviposition site choice and assessed the involvement of sensory organs in its detection. In two-choice oviposition assays, water conditioned with first-instar larvae attracted gravid females (OAI = 0.56), whereas water from fourth-instar larvae was repellent (OAI = -0.20), consistent with avoidance of suboptimal, resource-depleted habitats. Indole elicited strong oviposition attraction across a broad concentration range (0.1-50 µM), with no clear dose-response relationship. Surgical ablation of antennae and maxillary palps did not abolish indole-mediated preference but significantly reduced behavioral variability, suggesting that these structures modulate, rather than solely mediate, indole detection. Reanalysis of transcriptomes of antennae, maxillary palps, and legs in *An. gambiae* and *An. coluzzii*, along with quantitative RT-PCR in *An. stephensi*, revealed the expression of chemosensory genes (including *Obp1, Obp13, Obp25, Obp71, Or2*, and *Or10*) in the legs, indicating a potential role for leg chemosensation in oviposition decisions. These findings underscore the complexity of chemoreception and chemoperception in mosquito habitat assessment.

## 1. Introduction

Oviposition site selection is a critical stage in the mosquito life cycle, profoundly influencing population dynamics, larval development, and vector-borne disease transmission. Gravid females assess potential breeding habitats with care, aiming to maximize offspring survival while minimizing risks such as predation, desiccation, and resource scarcity. This decision-making process is shaped by a variety of environmental and sensory signals [1,2]. These oviposition decisions play a key role in determining the spatial and temporal distribution of mosquito populations, where even minor environmental characteristics can impact species persistence and pathogen transmission risk. Underpinning this complex behavior is an intricate sensory system, comprising the eyes, antennae, maxillary palps, and legs, that detects and integrates diverse environmental information [3,4]. The antennae function as the principal olfactory organ, detecting volatile chemical cues over long distances to guide females toward suitable oviposition sites. The maxillary palps assist in short-range odor and taste evaluation, while the legs facilitate contact chemosensation, allowing assessment of the site’s chemical makeup upon landing. Through multisensory integration, mosquitoes achieve refined discrimination among potential habitats, thereby enhancing the probability of successful egg deposition. Despite its importance, the understanding of the functional role of specific sensory appendages in oviposition site selection remains limited.

Humidity detection is critical for gravid mosquitoes when locating aquatic habitats to lay eggs. In *Anopheles gambiae* and *Aedes aegypti*, the ionotropic receptor gene *ir92a*, expressed in the antennae, plays a central role in mediating hygrosensation-driven water localization [5]. Chemical signals also significantly influence oviposition choices [6]. Volatile organic compounds such as indole, nonane, geosmin, fatty acids, and microbial metabolites function as attractants or deterrents, providing information about water quality, larval density, and the presence of conspecifics or predators [6]. Among microbial volatiles, indole (an aromatic heterocyclic compound) is particularly notable for its potent biological activity. Along with other volatiles, it guides gravid females in identifying suitable oviposition sites [7,8]. Indole triggers strong electrophysiological responses in the antennal sensilla of *An. gambiae* [7,9]. It is detected by odorant binding protein 1 (Obp1) [10], a process that may involve heterodimerization with Obp4 [11]. Odorant receptors Or2 and Or10 appear to function as specific receptors for indole and skatole, respectively [12,13], and share a common ancestral origin across mosquito lineages [14]. Although indole is commonly found in natural oviposition substrates [7,8,15], studies examining its role as a standalone compound in oviposition site selection remain limited.

Over the past decade, the originally Asian malaria vector mosquito, *Anopheles stephensi*, has invaded and established in Africa [16]. Its high vector competence for transmitting malaria presents a significant epidemiological concern [17,18]. The chemosensory gene repertoire of this species has been annotated at the genomic level [19,20]. However, oviposition behavior and the associated sensory mechanisms in *An. stephensi* remain poorly understood. Here, we demonstrate that indole acts as an oviposition attractant for gravid *An. stephensi* mosquitoes. Remarkably, the ablation of antennae and maxillary palps does not impair the indole-induced preference.

## 2. Methods and Materials

### 2.1. Mosquitoes

*An. stephensi* STE2 was acquired from the BEI MR4 repository. The mosquitoes were reared under controlled environmental conditions at 28°C and 80% relative humidity, with a 12-hour light/12-hour dark cycle. Blood feeding from mice was used to stimulate egg production. Adult mosquitoes were provided with a 10% sucrose solution. Eggs were transferred to water pans (20 cm × 30 cm × 3.5 cm) containing larval diet composed of brewer’s yeast and rodent food pellets in a 1:1 weight ratio.

### 2.2. Oviposition Activity Index Assay

Female mosquitoes aged 3-5 days were blood-fed on mice. Seventy-two hours after feeding, 60–70 gravid mosquitoes were released into a cage measuring 100 cm (L) × 50 cm (W) × 50 cm (H) for an oviposition bioassay. Two egg-collecting cups were positioned diagonally within the cage, each containing 200 mL of either control water or substrate water along with a piece of filter paper. The cups were placed approximately 110 cm apart. To minimize positional bias, the locations of the cups were switched between replicate experiments. The cups remained in the cage overnight to allow egg laying. Eggs deposited on the filter paper were subsequently counted under a microscope. Oviposition substrate preference was assessed using the Oviposition Activity Index (OAI), calculated as OAI = (Ns – Nc) /(Ns + Nc), where Ns represents the number of eggs laid in the substrate cup and Nc the number in the control cup. Substrates with an OAI value greater than zero were considered attractive, whereas substrates with an OAI value less than zero were considered deterrent.

### 2.3. Oviposition Response to Larval Water

The oviposition response of gravid mosquitoes to different habitat waters was evaluated using the OAI assay. Deionized (DI) water served as the control. The test substrates consisted of habitat water collected from rearing pans containing either 200 first- or fourth-instar larvae. Water containing first-instar larvae was collected 72 hours after egg hatching, while water with fourth-instar larvae was obtained when the larvae had reached the fourth instar stage. Prior to the assays, larvae were removed, and only the habitat water was used as the substrate.

### 2.3. Oviposition Response to Indole

Indole was obtained from Sigma-Aldrich. In the OAI assays, oviposition responses were evaluated with freshly prepared indole water at concentrations of 0.5, 1, 10, 30, and 50 μM, with DI water as the control.

### 2.4. Ablation of Antennae and Maxillary Palps in OAI Assays

We performed surgical ablation of the antennae and maxillary palps (AMP) in gravid mosquitoes. The AMP-ablated individuals were subsequently evaluated using an oviposition attraction index (OAI) assay to assess the role of these sensory structures in oviposition behavior. At 48 hours post-bloodmeal (PBM), females underwent AMP ablation and were maintained on 10% sucrose solution until 72 hours PBM, at which point their response to 1 µM indole was tested in the OAI assay. Oviposition behavior was compared between AMP-ablated mosquitoes and intact gravid controls. Both groups were derived from the same rearing cage, and OAI assays were conducted concurrently to reduce potential bias.

### 2.6. Statistical Analysis of OAI Data

Normality of OAI values was evaluated using the Shapiro–Wilk test. In all OAI assays, the data followed a normal distribution. One-sample t-tests, ANOVA, or unpaired t-tests were used as detailed in the main text. Comparisons of OAI between groups with ablated antennae and maxillary palps and intact controls were conducted using Welch’s t-test, while differences in variance between ablated and intact controls were assessed using the Levene test [21] and the Fligner–Killeen test [22]. All statistical analyses were performed at a significance level of α = 0.05.

### 2.7. RNA Extraction and Quantitative RT-PCR

Total RNA was isolated from antennae, maxillary palps, and legs (comprising the coxa, trochanter, femur, tibia, and tarsus). Tissue samples were homogenized in TRIzol reagent (Invitrogen), followed by RNA extraction and purification using the Invitrogen PureLink™ RNA Mini Kit in accordance with the manufacturer’s instructions. Genomic DNA was removed by Turbo DNase (Invitrogen) treatment. Total RNA concentration was quantified using a Qubit™ 4 Fluorometer (Thermo Fisher Scientific). For cDNA synthesis, more than 100 ng of total RNA was used as input. Reverse transcription was performed using the SuperScript™ IV kit (Invitrogen) according to the manufacturer’s protocol. Quantitative PCR was performed using Luna Universal qPCR Master Mix (New England Biolabs) on a CFX thermocycler (Bio-Rad), and data were analyzed with CFX Maestro software (Bio-Rad). In qRT-PCR data analysis, target gene transcript levels were normalized to the level of the gene encoding the 40S ribosomal protein S29, which was selected for its even expression across the sensory tissues examined in this study. Primer sequences are provided in Table 1.

**Table 1.**
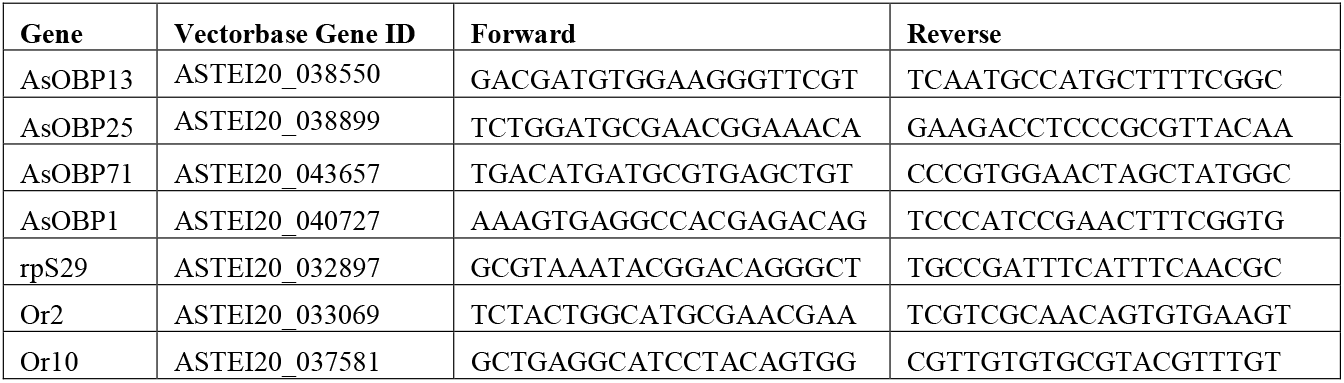
PCR Primers.

### 2.8. Reanalysis of RNA-seq Datasets

Three previously published RNA-seq datasets profiling the chemosensory transcriptomes of antennae, maxillary palps, and legs [23-25] (Table 2) were reanalyzed using the RNA-seq analysis module of the CLC Genomics Workbench v.25 (Qiagen). *Anopheles coluzzii* [26], originally known as the M molecular form of *An. gambiae* [27]. The protein-coding genes have nearly identical sequences between *An. gambiae* and *An. coluzzii*. Therefore, RNA-seq reads from both species were aligned to the *An. gambiae* PEST genome reference. Briefly, raw reads were trimmed and then mapped to annotated genes from the PEST genome (VectorBase release 68). Transcripts per million (TPM) values were compared across conditions as described in th original studies.

**Table 2.** RNA-seq datasets of chemosensory organs.

| Species | Organ | BioProject | PMID | Reference |
| --- | --- | --- | --- | --- |
| *An. gambiae* | Antenna | PRJNA215079 | 23630291 | [24] |
| *An. coluzzii* | Leg | PRJNA910489 | 34903168 | [23] |
| *An. coluzzii* | Antenna, maxillary palp | PRJNA400609 | 28938869 | [25] |

## 3. Results

### 3.1. Oviposition Response to Habitat Water

We employed the OAI assay to evaluate the oviposition response of gravid mosquitoes to different substrates. DI water served as the control. Under conditions with two control water cups present in the assay cage, gravid mosquitoes were expected to exhibit no preference. Consistent with this expectation, the OAI values were randomly distributed across both sides over 11 experimental replicates. The mean OAI was -0.04 (95% CI: –0.22 to 0.14; one-sample t-test, t = 0.488, p = 0.629), suggesting no inherent bias in the experimental setup and indicating that mosquitoes detected and oviposited equally in both water containers (Figure 1A). Subsequently, we assessed oviposition preferences for larval water conditioned with first- and fourth-instar larvae. As illustrated in Figure 1B, water from the first-instar stage was attractive to gravid mosquitoes, yielding an OAI of 0.56 ± 0.07. In contrast, water from the fourth-instar stage elicited a deterrent effect, with an OAI of -0.20 ± 0.09. An unpaired t-test revealed a statistically significant difference in preference (t = 6.543, p < 0.0001). The estimated OAI difference between the two substrates was 0.76 (95% CI: 0.50 to 1.02), supporting a strong substrate-specific effect on oviposition behavior.

**Figure 1.**
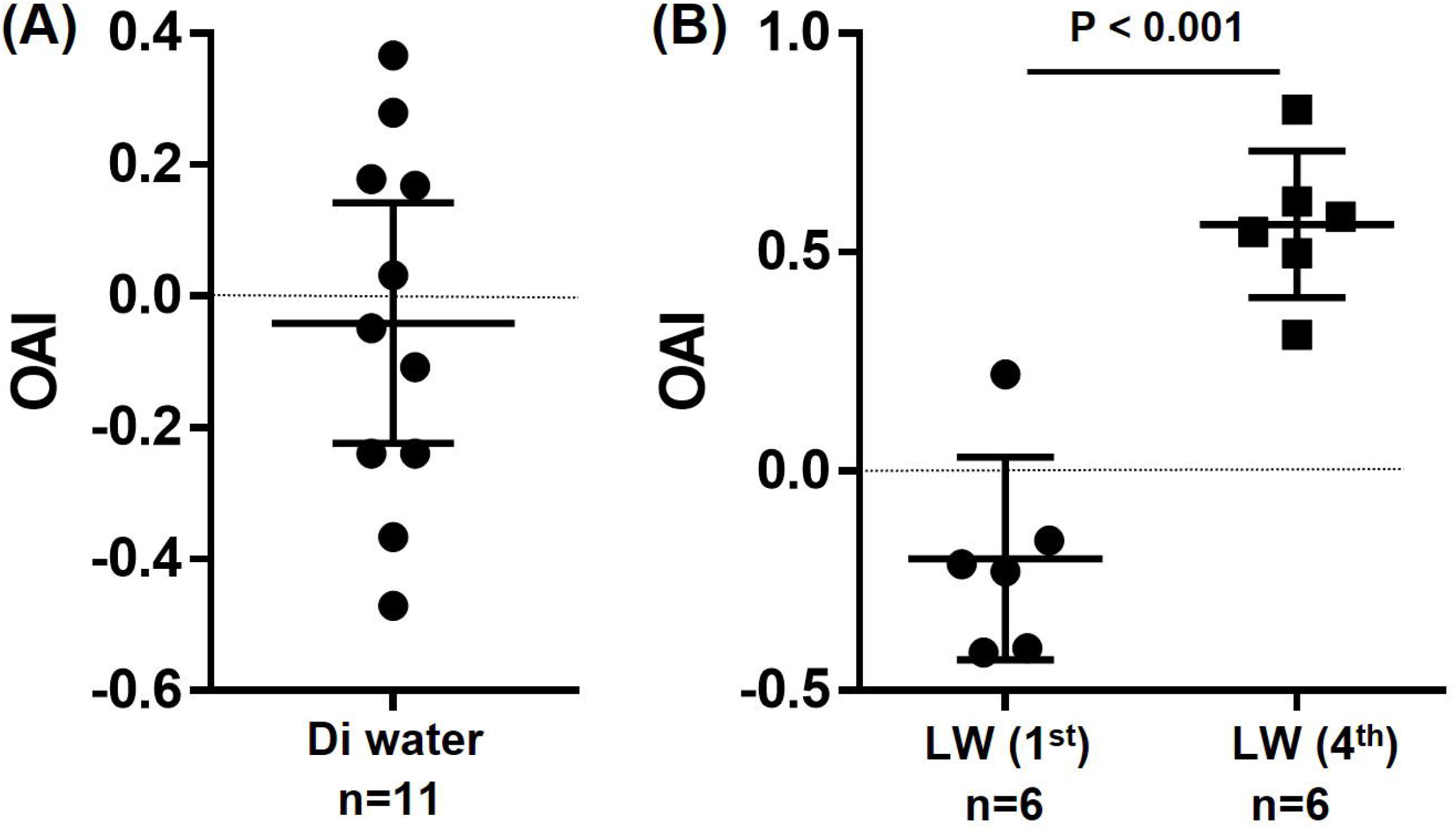
Oviposition preference assays using habitat water. Each data point represents the OAI from a single replicate. Mean OAI values are shown with 95% confidence intervals. (A) In assays containing two cups of DI control water, OAI values were randomly distributed, indicating no preference (one-sample t-test, t = 0.488, p = 0.629). (B) OAI values for larval water (LW) collected at the first and fourth instar stages differed significantly (unpaired t-test, t = 6.543, p < 0.0001).

### 3.2 Oviposition Response to Indole

To assess the influence of indole on oviposition preference in gravid mosquitoes, OAI was measured at six indole concentrations (0.1, 0.5, 1, 10, 30, and 50 μM), with DI water as the control. Gravid mosquitoes consistently showed positive OAI values, indicating a preference for indole-containing substrates over the control. No dose-dependent effect on oviposition preference was detected across the tested concentrations (one-way ANOVA, F = 0.610, p = 0.692).

### 3.3. Effect of Antennal and Maxillary Palp Ablation on Oviposition Response

Obp1 and Obp4, which are indole-responsive [10,28], are expressed abundantly in the antennae and maxillary palps in *An. gambiae* [24] and *An. coluzzii* [25]. To assess the role of these sensory organs in mediating indole preference during oviposition, we surgically removed the antennae and maxillary palps from blood-fed mosquitoes at 48 hours PBM. We then evaluated the OAI using 1 μM indole in both ablated and control gravid mosquitoes. As shown in Figure 2B, no significant OAI difference was observed between ablated and intact control mosquitoes (Welch’s t-test, t = 2.097, p = 0.061), suggesting that antennae and maxillary palps are not essential for indole detection under these experimental conditions. Intriguingly, OAI values in ablated mosquitoes exhibited less variability than those in intact controls (Figure 2B). The mean OAI was numerically higher in ablated females (M ± SD = 0.907 ± 0.027) than in intact females (M = 0.750 ± 0.087), indicating greater response variability among intact individuals. The heterogeneity in variance between the two groups was confirmed by an F-test (F = 12.440, p = 0.009), as well as by Levene’s test (L = 6.889, p = 0.017) and the Fligner-Killeen test (FK = 5.309, p = 0.021). The markedly narrower distribution of OAI values in the ablated group suggests greater behavioral consistency. Overall, the OAI distribution patterns in Figure 2B indicate that ablation of antennae and maxillary palps did not diminish the overall attraction to indole.

To determine the role of antennae and maxillary palps in the indole preference in the oviposition, we surgically ablated the antennae and maxillary palps from the blood-fed mosquitoes at 48 hr PBM. Then we tested the OAI with 1 μM indole in ablated and control gravid mosquitoes. As shown in Figure 2B, the OAI between the ablated and the intact control mosquitoes was not significantly different, as tested by the Welch’s t-test (t=2.097, p=0.061), suggesting that antennae and maxillary palps do not play a necessary role for the indole detection in the assay. Intriguingly, the OAIs of ablated mosquitoes clustered more closely than those of intact controls (Figure 2B). The ablated females showed a numerically higher mean (M ± sd = 0.907 ± 0.027) than intact females (M = 0.750 ± 0.087), suggesting that the responses were more variable in the intact mosquitoes. The unequal OAI variances between them were confirmed by the F test (F = 12.440, p = 0.009) and two additional variance tests: the Levene test (L = 6.889, p = 0.017) and the Flinger-Killeen test (FK = 5.309, p = 0.021). The significantly narrower range of OAI values in the ablated group indicates greater behavioral consistency. Overall, the OAI distribution patterns shown in Figure 2B suggest that ablation of the antennae and palps did not diminish overall attraction to indole; however, it likely reduced exposure to competing sensory noise, which, in intact females, contributes to greater response variability.

**Figure 2.**
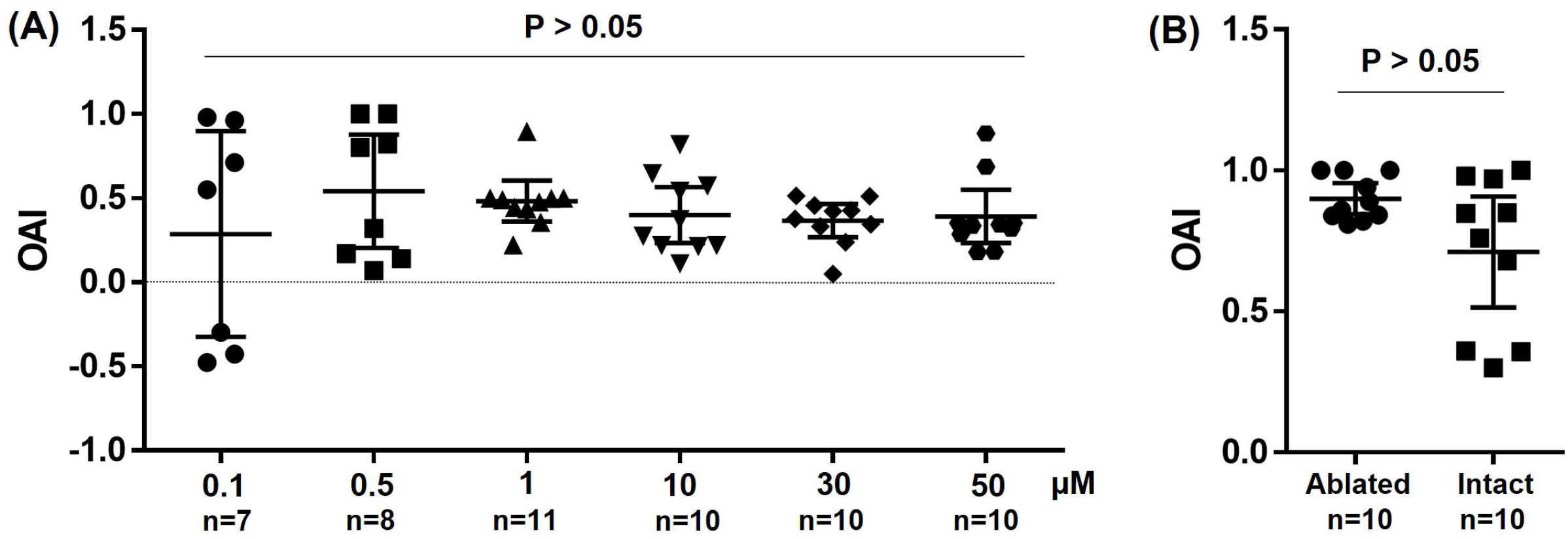
Oviposition preference for indole. Each data point corresponds to the OAI from one replicate, with the number of replicates (n) indicated. (**A**) Dose-response assays for indole at various concentrations. No significant differences in OAI were observed across the tested concentrations (one-way ANOVA, F = 0.610, p = 0.692). (**B**) Oviposition assays comparing mosquitoes with and without antennae/maxillary palps at 1 µM indole. The OAI did not differ significantly between groups (Welch’s t-test, t = 2.097, p = 0.061). However, the variances in OAI were unequal (Levene’s test, L = 6.89, p = 0.017; Fligner-Killeen test, FK = 5.31, p = 0.021).

### 3.4. Chemosensory Transcriptome Data Mining

The findings from the above oviposition assays indicate that sensory structures beyond the antennae and maxillary palps may be involved in detecting indole during oviposition. However, the role of the legs in habitat selection is poorly understood. It has been proposed that tarsal sensilla may perceive taste signals upon contact with the water surface of the habitat [2,29]. Several chemosensory transcriptome datasets are available from previous studies, including those of *An. gambiae* antennae [24], *An. coluzzii* antennae, maxillary palps [25], and legs [23]. To compare transcript abundance of sensory genes encoding odorant-binding proteins, odorant receptors, gustatory receptors, and ionotropic receptors across these tissues, we reanalyzed these datasets. Since the datasets were generated in different studies, direct comparison of expression levels may not be appropriate. We therefore normalized the TPM values for each gene by dividing them by the TPM of BTF3 (AGAP006614), which encodes basic transcription factor 3. BTF3 is a conserved transcriptional regulator that acts as a general cofactor for RNA polymerase II [30], indicating its involvement in gene transcription and its suitability as an indicator of general transcriptional activity. Across the datasets, BTF3 exhibited a TPM range of 307–732 (Table S1), which is lower than that of the gene encoding 40S ribosomal protein S7 (AGAP010592; TPM range: 893–3250), a commonly used housekeeping gene in qPCR normalization. After normalization to BTF3 TPM, the maximum normalized TPM value for each gene was scaled to 100, and all other TPM values were divided by this maximum to represent relative abundance across organs. Figure 3 displays a heatmap of relative transcriptional abundances for 92 genes whose original, unnormalized TPM values exceeded 20 in at least one organ. The sensory genes were grouped into three clusters based on their relative abundance: genes with higher expression in antennae, palps, and legs, respectively (Figure 3A). These genes showed organ-specific expression patterns. For instance, *Obp20, Obp47, Obp3* (AGAP001409), *Obp7, Obp66, Obp1, Obp4*, and *Obp5* were predominantly abundant in antennae; *Obp48* was abundant in both antennae and palps; *Obp57* was abundant in palps and legs. In contrast, *Obp13, Obp71, Obp49*, and *Obp54* were primarily abundant in legs (Figure 3B). Obp genes were expressed at substantially higher levels than Or, Gr, and Ir genes. *Or2* and *Or10* were abundantly expressed in antennae and palps (Figure 3C).

**Figure 3.**
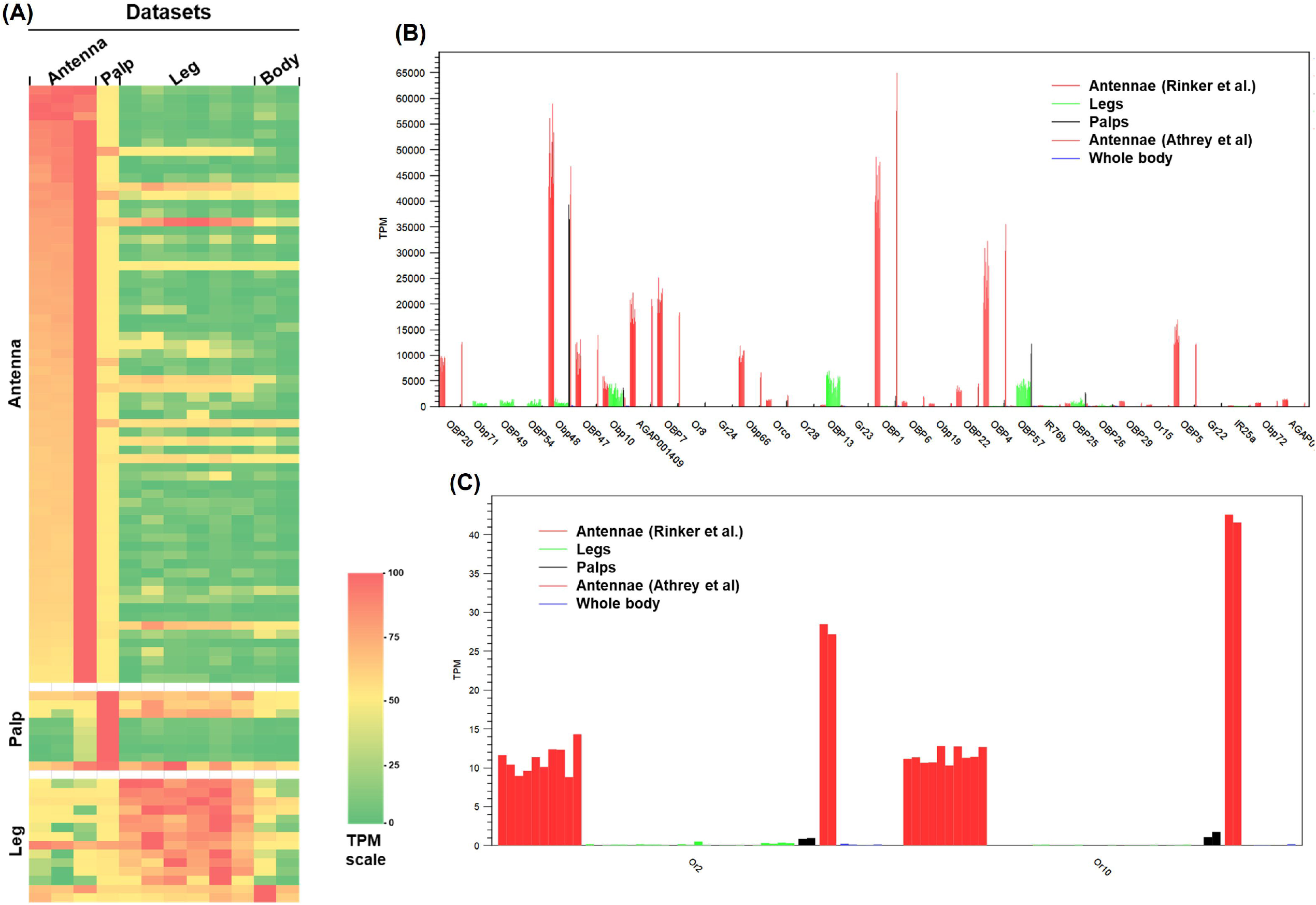
Relative transcriptional abundances of chemosensory genes in different sensory organs in *An. gambiae* and *An. coluzzii*. **(A)** The heatmap illustrates the relative abundance of sensory genes (Obp, Or, Gr, and Ir) with TPM values exceeding 20 in at least one dataset. Three gene clusters, each enriched with genes predominantly expressed in the antenna, maxillary palp, and leg, are labeled on the left side. Gene names, original TPM values, and gene function annotation are provided in Table S1. The three antenna datasets include sugar-fed (pooled) and blood-fed (pooled) samples of *An. gambiae* [24], as well as sugar-fed antennae of *An. coluzzii* [25]. **(B, C)** TPM profiles of representative genes showing differential expression across replicates of sensory organs. Organs are color-coded for display.

In *An. stephensi*, the expression profiles of sensory genes have not yet been characterized at the transcriptome level. Using qRT-PCR, we compared the transcript levels of *Obp1, Obp13, Obp25, Obp71, Or2*, and *Or10* across antennae, palps, and legs. As illustrated in Figure 4, the expression levels and reproducibility among three biological replicates for each chemosensory gene are visually summarized. *Obp1* was highly expressed in antennae and legs relative to palps, whereas *Obp13, Obp25*, and *Obp71* showed greater abundance in legs than in antennae or palps. *Or2* and *Or10* belong to the conserved indole-sensitive *Or2/Or10* clade [14]. *Or2* was detected in all three organs, with considerable variation observed among replicates within each tissue. In contrast, *Or10* expression was markedly higher in legs compared to antennae and palps.

**Figure 4.**
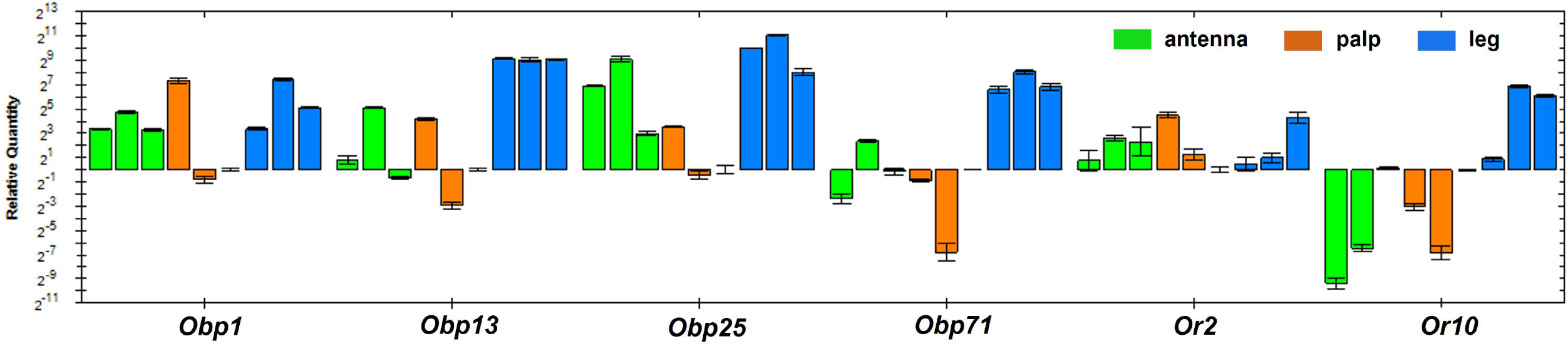
Relative abundance of selected genes in the antennae, palps, and legs measured by qRT-PCR. Three biological replicates were performed for each tissue. Expression levels were normalized to the sample from palp replicate 3. Error bars indicate the variation between two technical replicates on the qPCR plate.

## 4. Discussion

Oviposition site selection is critical in the mosquito life cycle. Substantial research has been conducted in *An. gambiae* to identify the cues that gravid mosquitoes detect during oviposition site selection. *An. stephensi*, originally a malaria vector native to South Asia and the Persian Gulf region, has emerged over the past decade as an invasive, urban-adapted vector that is now well-established and expanding across numerous African countries [31]. The ecology of invasive *An. stephensi* in Africa has been the subject of recent studies [16]. Nevertheless, knowledge regarding the chemical ecology governing its oviposition behavior remains limited [6,32]. In *An. coluzzii*, experimental evidence indicates that water conditioned by low-density or early-instar larvae attracts gravid females, whereas water conditioned by high densities and/or later instars (including fourth instars) becomes increasingly repellent. The oviposition index shifts from positive to strongly negative as larval developmental stage and conditioning duration increase [33]. In this study, we found a similar trend in *An. stephensi* (Figure 1). Gravid females deposit significantly more eggs in cups containing water previously inhabited by first-instar larvae compared to water from fourth-instar larvae, indicating an attraction to cues from early larval stages and a deterrent effect from those of later stages. This attraction to first-instar water is likely due to lower concentrations of larval and microbial metabolites, suggesting reduced competition for resources. In contrast, fourth-instar water may reflect resource depletion, increased oxygen and organic matter consumption. According to Suh et al. (2013), repellent volatile semiochemicals, such as dimethyl disulfide (DMDS), dimethyl trisulfide (DMTS), and 6-methyl-5-hepten-2-one (sulcatone), were detected in water from suboptimal larval habitats [33]. In a pull-push assay, *An. gambiae* s.l. was attracted to nonane and 2,4-pentanedione (2,4-PD), both found in larval water across stages, but repelled by DMDS and DMTS, which are present predominantly in late-stage larval water [34]. In our experimental setup, metabolites derived from mosquito larvae and associated microorganisms accumulate as larvae develop. Further research is needed to identify the specific chemical compounds responsible for these attractive and repellent effects.

Indole and its derivatives are recognized as key infochemicals that, depending on the context, guide mosquitoes in locating and assessing aquatic oviposition sites. Their functions vary by species, concentration, and the surrounding chemical mixture. Both indole and 3-methylindole (skatole) have been identified as volatile compounds associated with oviposition sites [7,15,35,36]. Previous research on chemically mediated oviposition has largely employed assays using water containing bacteria or plant-derived blends [8,37]. In the present study, we examined the influence of indole on oviposition behavior to assess its contribution to habitat selection. Indole-containing water attracted mosquitoes to oviposit across a concentration range of 0.1-50 µM. The effect did not appear to be dose-dependent, with a marginally effective dose at 0.1 µM (Figure 2A). It has been reported that Obp1 in *An. gambiae* binds indole with high affinity, exhibiting an equilibrium dissociation constant Kd of 2.3 µM [10]. In our experimental setup, attractive indole concentrations were as low as 0.1 µM, suggesting that *An. stephensi* indole-binding Obps have a high affinity to indole volatiles. Furthermore, the experimental setup allowed landing on filter paper in indole cups, enabling near proximity to the compound and potentially enhancing cue detection.

The most striking finding of this study is that surgical removal of the antennae and maxillary palps did not attenuate the oviposition preference mediated by indole (Figure 2B). Although the mean OAI did not differ significantly between ablated and intact mosquitoes, the ablated group showed a significantly lower index variance. This lower variability suggests that inputs from the antennae and maxillary palps may introduce modulatory or competing sensory signals, thereby increasing behavioral variability rather than being essential for indole detection. The persistent attraction to indole even after removal of the antennae and palps suggests the involvement of additional chemosensory structures, challenging the prevailing view that antennae and maxillary palps are the primary olfactory organs guiding oviposition site selection.

Legs emerge as potential candidates for indole detection in this context. Transcriptome reanalysis and qRT-PCR experiments demonstrate that several chemosensory genes, including *Obp1, Obp13, Obp25, Obp71, Or2*, and *Or10*, are abundantly or preferentially expressed in the legs (Figures 3 and 4). Notably, Or2 and Or10 belong to an evolutionarily conserved clade known to mediate the detection of indole and skatole across Dipteran lineages [14,38]. The detectable expression of *Or2* and *Or10* in the legs, together with the other leg-enriched Obp genes, raises the possibility that tarsal olfactory chemoreception may contribute functionally to habitat evaluation. In the OAI assays, gravid females land on filter paper, bringing their tarsi into near proximity with the indole-containing water surface. Under these experimental conditions, tarsal chemosensory input may complement or even dominate volatile detection, suggesting a multi-organ model for indole detection in *An. stephensi*. These findings warrant further investigation to test the hypothesis that oviposition site selection involves multiple sensory organs acting redundantly or synergistically in detecting olfactory cues. While antennal detection of long-range volatiles may guide initial orientation toward breeding sites, close-range or contact-mediated detection by tarsi may finalize the oviposition decision. The persistence of indole preference after antennal ablation indicates that tarsal chemosensation alone is sufficient under these laboratory conditions. Whether this holds in natural settings with greater spatial complexity and competing cues remains to be tested.

In *Culex quinquefasciatus*, the labrum within the proboscis contributes to short-range detection of 4-ethylphenol (4EP), a compound that modulates both oviposition and blood-feeding behaviors [39]. Notably, the indole receptor *Or10* ortholog *CqOr21* was also found to be expressed in the proboscis stylet, although the functional role of the proboscis in responding to indole was not examined in that study [39]. In our experimental setup, the proboscis remained intact. Thus, we cannot exclude the possibility that the proboscis participates in indole reception under our conditions.

Overall, this study demonstrates that oviposition chemosensation in *An. Stephensi* involves a system with multiple sensory appendages, suggesting that the legs may play a previously underappreciated role in habitat selection behavior. From a vector control perspective, clarifying the chemosensory mechanisms underlying oviposition opens avenues for developing attract-and-kill or push-pull strategies targeting gravid mosquitoes [4]. The ongoing invasive spread of *An. stephensi* throughout Africa underscores the urgency for interventions tailored to species-specific or behavioral traits. Field validation of indole-based oviposition attractants may offer crucial translational insights.

## Supplementary Materials

Table S1: Scaled TPM heatmap

## Author Contributions

Conceptualization, J.X.; methodology, J. A., J.X.; formal analysis, J. A., J.X.; writing—original draft preparation, J.A.; writing—editing and finalizing, J. X.. All authors have read and agreed to the published version of the manuscript.

## Data Availability Statement

This study used three RNA-seq datasets previously published by other research teams. The BioProject IDs for these datasets are provided in Table 2.

## Acknowledgments

The authors thank the Biology Department, College of Arts and Sciences, New Mexico State University, for the financial support provided. During the preparation of this manuscript, the authors used Perplexity.ai for organizing the relevant literature for the Introduction and Discussion sections. Grammarly was integrated into MS Word while writing to refine grammar, tone, and clarity. The authors have reviewed and edited the output and take full responsibility for the content of this publication.

